# Demographic, behavioral, and ecological data from a long-term field study of wild baboons in Amboseli, Kenya

**DOI:** 10.1101/2025.10.03.680086

**Authors:** Chelsea A. Southworth, Jack C. Winans, Jacob B. Gordon, Niki H. Learn, William A. Wilber, Catherine Andreadis, Gretchen Andreasen, Mimi Arandjelovic, C. Ryan Campbell, Mary N. Chege, Maria J. A. Creighton, Carmen M. Cromer, Reena Debray, Carly C. Dickson, Pamela Ferretti, Elizabeth M. George, Laurence R. Gesquiere, Shuyu He, Leif Hey, Emily E. Jefferson, Ipek G. Kulahci, Brian A. Lerch, Lee Nonnamaker, Iker Rivas-González, Beniamino Tuliozi, Shasta E. Webb, Susan C. Alberts, Elizabeth A. Archie, Jenny Tung

## Abstract

Long-term data sets on individually recognized animals and their environments are critical to understanding animal behavior, evolution, and ecology. However, they are resource- and time-intensive and seldom made publicly available. The Amboseli Baboon Research Project (ABRP) is one of the longest-running studies of a wild mammal population in the world and has collected extensive data on the baboon population of the Amboseli ecosystem in Kenya since 1971. Here, we describe four ABRP data sets newly available to the evolutionary biology, behavioral ecology, and primatology communities: (1) the sizes and demographic compositions of 21 social groups from 1971-2023; (2) the activity budgets of adult females and immatures from 1984-2023; (3) behavioral data on diet for adult females and immatures from 1984-2023; and (4) weather data, including precipitation from 1976-2023 and temperature from 1976-2022. Data are aggregated annually and monthly to enable cross-data set analyses. These data offer a rare longitudinal perspective on behavioral and ecological change in a wild mammal population.

## Background

Longitudinal data on wild animal populations are essential to understanding the relationships between animal behavior, ecology, and fitness. Amassing long-term data sets that span the lives of many animals allows researchers to investigate these relationships with considerable power^1,2^. However, life course data of this scale are rare for long-lived species^3^, even though cross-sectional data often fail to capture the full range of social and environmental conditions that individuals experience^1^. To spur methods development, facilitate comparative analyses, and understand change over time, long-term data sets on relevant behavioral, demographic, and environmental variables are required, and, where feasible, should be made publicly available.

The Amboseli Baboon Research Project (ABRP) is one of the longest-running field studies of a wild mammal population in the world. Founded in 1971 by Jeanne and Stuart Altmann, the ABRP has observed the baboon population living in the Amboseli basin for more than 50 years^4–6^. Baboons are highly social, ecologically flexible primates that are widely distributed across Africa and the Arabian peninsula^7^. In Amboseli, baboons live in stable social groups of varying sizes (∼20 to over 120 individuals), consisting of multiple adult females and males as well as immatures of both sexes^6^. Females are philopatric, while males typically disperse to other social groups at sexual maturity and can move between groups multiple times in their lives^8–12^. The baboons in this ecosystem, like many baboon populations across Africa, experience highly variable environments, including pronounced seasonal variation and high yearly variance in temperature and rainfall conditions, even within season^13,14^. They cope with this variability through behavioral flexibility—adjusting activity budgets and capitalizing on seasonal food availability as highly selective ‘eclectic omnivores’^7,13,15–17^. These ecological and dietary pressures in turn shape many aspects of the baboons’ lives, including their maturation and reproductive schedules, body condition, and movement patterns^13,15,17–21^.

The ABRP conducts intensive data collection on multiple social groups with overlapping home ranges, alongside opportunistic monitoring of other groups that range in the study area^4,5^. Over the course of the project, more than 2,000 individually-recognized animals have been monitored^6^. Here, we describe and make newly available four of the resulting long-term data sets: 1) group demography data for 21 social groups that were followed for different periods over 52 years, 2) activity budgets of adult females and immatures of both sexes over 39 years, 3) diet composition at the social group and study population levels over 39 years, and 4) rainfall and temperature measurements over 47 years. Each data set is aggregated over the same timescales (months and years), facilitating cross-data set analyses. We provide visual summaries of patterns across time and social groups for each dataset, as well as cleaned, aggregated, validated, and processed data for other researchers to incorporate in their own analyses. We hope that these data will facilitate comparative analyses of animal behavior and behavioral and ecological change over time and may be useful as a teaching tool. In addition, this repository will be useful for the many researchers working with ABRP data or other east African mammals as an open-access reference for demographic, behavioral, and ecological patterns in this ecosystem, from 1971 to 2023.

## Methods

### Study site and subjects

Since 1971, the ABRP has studied a naturally admixed population of yellow (*Papio cynocephalus*) and anubis baboons (*P. anubis*; most population members are of predominantly *P. cynocephalus* ancestry^22,23^) located in the Amboseli ecosystem of southern Kenya (2°40’S, 37°15’E, 1100 m altitude)^5^. Amboseli is a semi-arid, short grass savannah ecosystem that experiences considerable seasonal and inter-annual variation in precipitation and temperature^13,14^ (Fig. 1). Generally, four seasons of varying lengths characterize intra-annual rainfall patterns in Amboseli^14^. The hydrological year begins with a short wet season lasting from November to January, followed by a short dry season in February, a long wet season from March to May, and a long dry season from May to October (Fig. 1B). In all data sets provided here, the hydrological year therefore runs from November 1 to October 31 (e.g., hydrological year 2023 encompasses November 1, 2022 to October 31, 2023). Rain almost never falls in the long dry season, but Amboseli experiences substantial inter-annual variability in rainfall during the rest of the hydrological year (Fig. 1D). Daily temperatures peak in February and March during the short dry and early long wet season and fall to their lowest in the long dry season from June to August (Fig. 1A). Maximum and minimum temperatures vary inter-annually, but are more consistent compared to rainfall (Fig. 1C).

**Figure 1.**
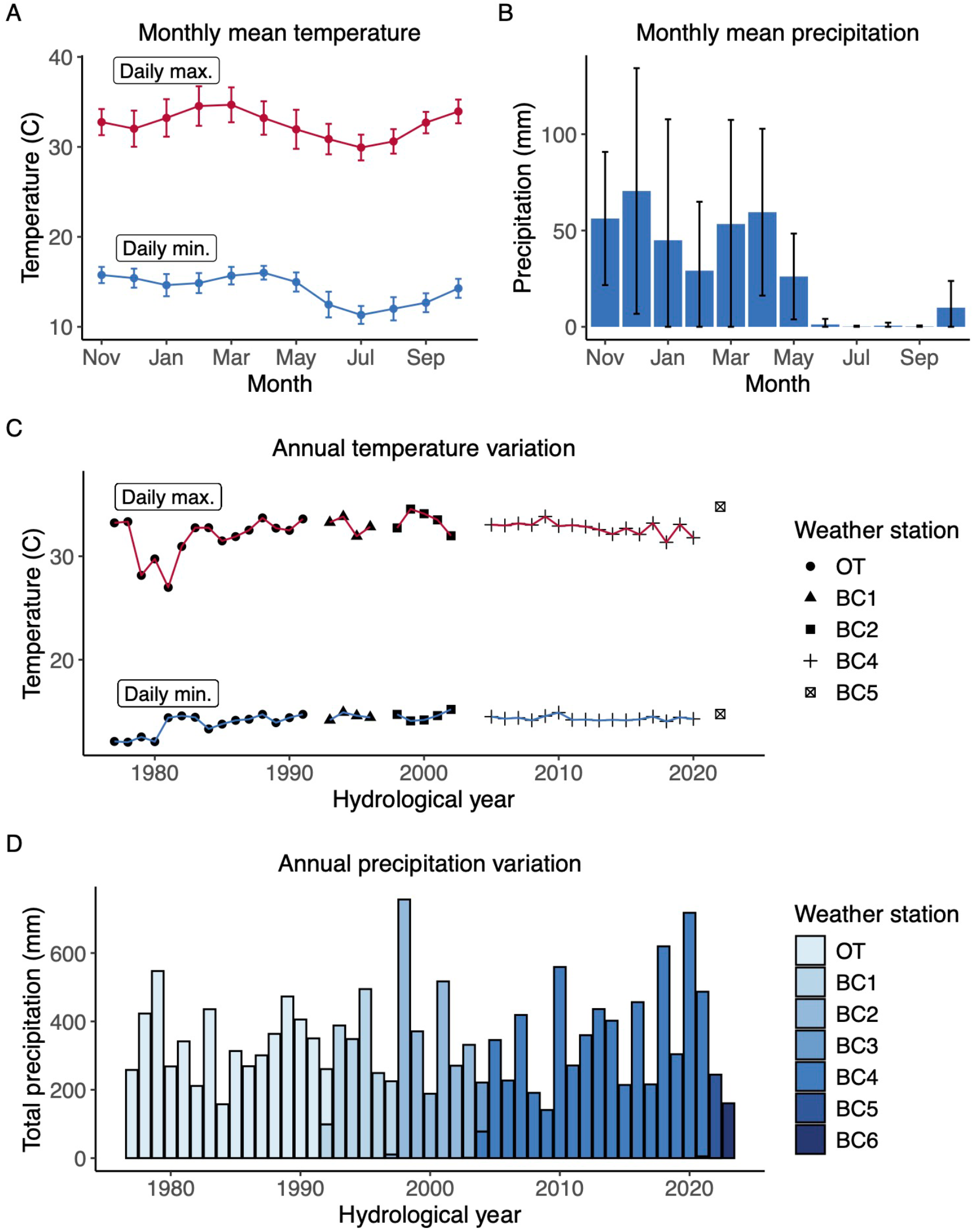
Monthly and annual patterns of temperature and precipitation in Amboseli, Kenya. Temperatures were taken from analog min-max thermometers and precipitation measures were taken from a standard rain gauge. The location of the thermometers and rain gauges changed slightly over the years but, with one exception, were within 150 m of each other. (A) and (B) depict monthly patterns over the hydrological year, which runs from November to October. In (A), points represent mean daily maximum (red) and minimum (blue) temperatures and error bars represent standard deviations within months between hydrological years 1977 and 2022. In (B), bar heights represent mean precipitation volume in each month and error bars represent standard deviations within months between hydrological years 1977 and 2023. (C) depicts annual average minimum and maximum temperature from hydrological years 1977 to 2022. (D) depicts total annual precipitation from hydrological years 1977 to 2023. In (C), the black points on the red and blue lines depict the mean maximum and minimum daily temperature, respectively, in each hydrological year. Annual mean temperature values were only included in (C) if all data for a given hydrological year were collected by the same weather station. In (D), the bars depict the total amount of rainfall in each hydrological year.

Over the course of the long-term study, the ABRP has studied between one and six social groups at a time, all descendants through fission or fusion of two original study groups on which observation began in 1971 and 1980, respectively^5^ (Fig. 2). Throughout the study period, data were consistently collected on demography, behavior, and reproductive patterns; collection of biological samples for genetic and hormone analysis began respectively in the 1980s and 1990s^5^. We distinguish between social groups that were regularly subject to such intensive data collection (‘study groups’) and other social groups that were present in the ecosystem but were monitored on an irregular or infrequent basis (‘non-study groups’). The number of study groups has varied over time due to the number of project observers, changes in the size of the population, and permanent fissions or fusions (Fig. 2). To maintain high levels of data collection effort across study groups, some study groups were dropped from intensive data collection over time (Fig. 2). Detailed descriptions of the data collection protocols maintained by ABRP can be found online at the ABRP’s website^24^.

**Figure 2.**
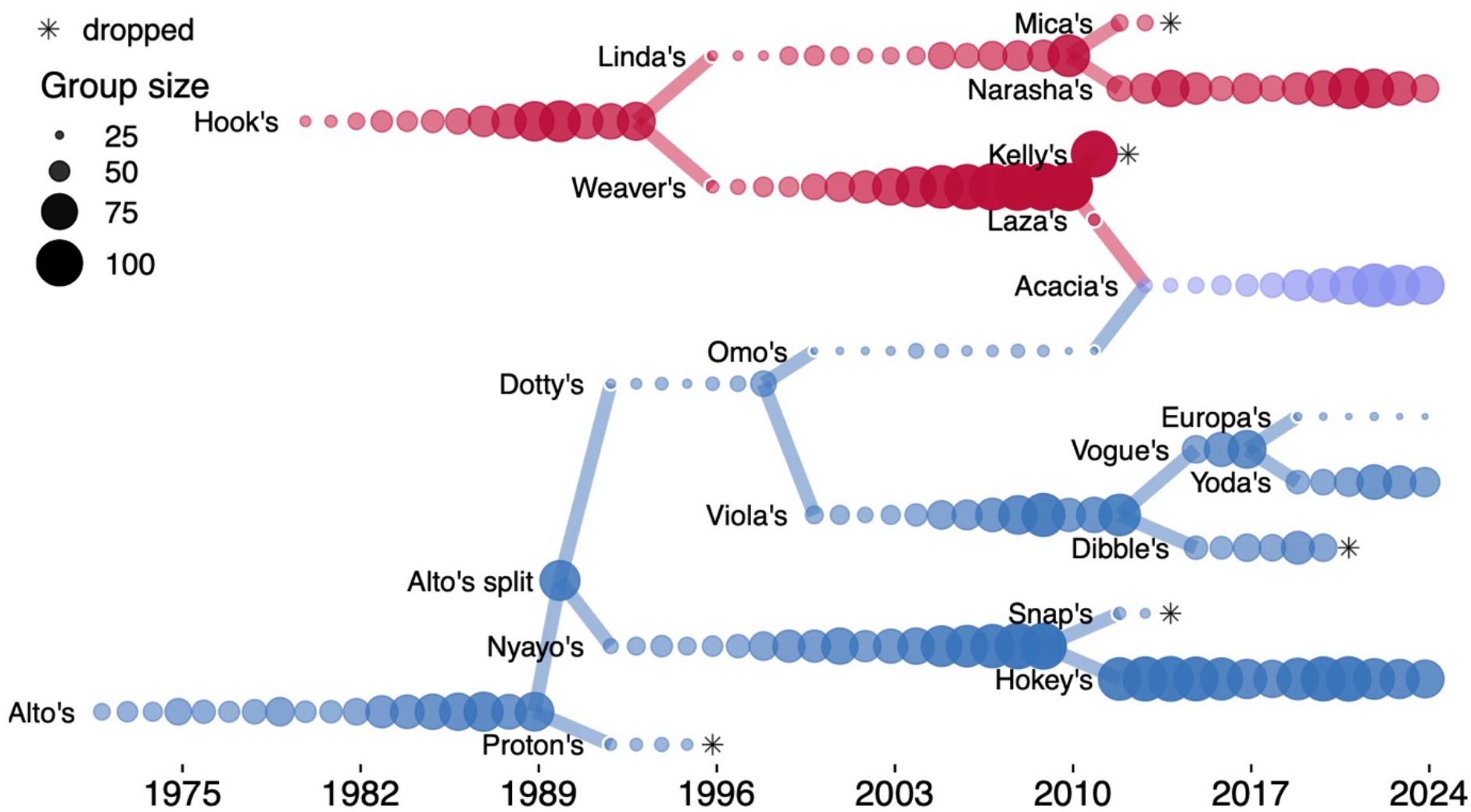
Total size of social groups subject to intensive monitoring by the Amboseli Baboon Research Project over time. Each group is represented by one data point for each hydrological year that it was intensively monitored and included in the study. Point size varies by group size, and lines connecting groups represent fission events (when groups permanently split) and fusion events (when groups permanently merged). Alto’s group, one of the two original study groups, and its daughter groups are represented in blue, and Hook’s group, the other original study group, and its daughter groups are represented in red. Acacia’s group was formed by the fusion of one daughter group from each lineage and is represented in purple. Asterisks indicate instances in which a group was dropped from intensive data collection.

### Demographic and behavioral data collection

The group demographic data were compiled from census records collected between 1971 and 2023, while the activity budget and diet data came from focal animal samples conducted between 1984 and 2023. Experienced field researchers performed group censuses and focal animal samples in the context of half-day visits to each group, which were conducted in either the morning or afternoon 6 days per week. At this schedule, the typical frequency with which each group was observed ranged from 2-4 days per week to near-daily, depending on the number of groups being monitored at a time and the number of project observers.

Census data were collected at the start of each group visit, including information on the number and identity of all individuals present in the group and on any demographic events, including births, known or suspected deaths, and known or suspected dispersal events. From these data, we calculated for each social group in each month and hydrological year: i) the total number of unique individuals that were resident in the group, including ii) the total number of unique adult females, iii) the total number of unique adult males, and iv) the total number of unique non-adult individuals. We used the concept of ‘residency’ to distinguish stable group membership from unstable ‘floating’ between groups, which was generally seen only in adult males but sometimes happened with other age-sex classes during group fissions. In principle, an individual was considered ‘resident’ in a given social group over a given period of time if they were present in the group more frequently than they were absent from it. In practice, residency was calculated algorithmically for each individual on each day via day-by-day assessments of their census observations (for full details, see^25^). For the yearly demographic data set, any female who had reached menarche before or during a given hydrological year was considered an adult in that year and any male who had attained a dominance rank above at least one other adult male before or during a given hydrological year was considered an adult in that year^26,27^. Analogously, for the monthly demographic data set, individuals who reached adulthood before or during a given month were counted as adults for that month.

These demographic data were collected from 21 unique social groups, including 185 complete group-years and 31 partial group-years, and 2,372 group-months. Partial group-years occurred when groups fissioned or fused, or when logistical obstacles interfered with intensive data collection on a group in a given year. When a group underwent a permanent fission or when two groups fused, the products of the fission/fusion were considered new social groups, i.e., no group retained an original group’s ‘identity.’

During group visits, focal animal samples were conducted on adult females and on juveniles of both sexes in a pre-determined, randomized order^28^. Focal animal samples and the resulting activity budgets and diet data were typically collected between 0700 and 1100 or between 1400 and 1800. Samples lasted 10 minutes and on each minute during the sample, the focal animal’s activity (feeding, walking, resting, or socializing) was recorded (Fig. 3). Beginning in September 1985, observers began additionally recording the type of food focal individuals were eating when the activity was ‘feeding’ (Fig. 4). For some food types (e.g., diet items from *Vachellia* trees), the species name and plant part were recorded, but for others (e.g., diet items from some grasses), species identification was not consistently possible. Therefore, we categorized diet items into eight types based on phylogenetic and/or structural properties that together make up 93.5% of total time spent feeding during years when food types were identified (see Fig. 4). We consider these eight types to vary meaningfully with respect to nutritional value, processing demands, spatio-temporal dispersion, and other characteristics of biological significance to foraging baboons^15^. Of these types, we consider grass corms, grass blade bases and seed heads, and *Vachellia* seeds to be low-energy foods, and we consider grass blades, fruits, flowers, tree gum, and invertebrates to be high-energy foods^14,15^. The diet items that constituted the remaining 6.5% of feeding time, including unidentifiable diet items, were classified into a ninth category called ‘Other’. Because observers did not record any information about food types during focal animal sampling from January 1984 through August 1985, all feeding observations from this period were labelled as occurring on ‘Unclassified’ food.

**Figure 3.**
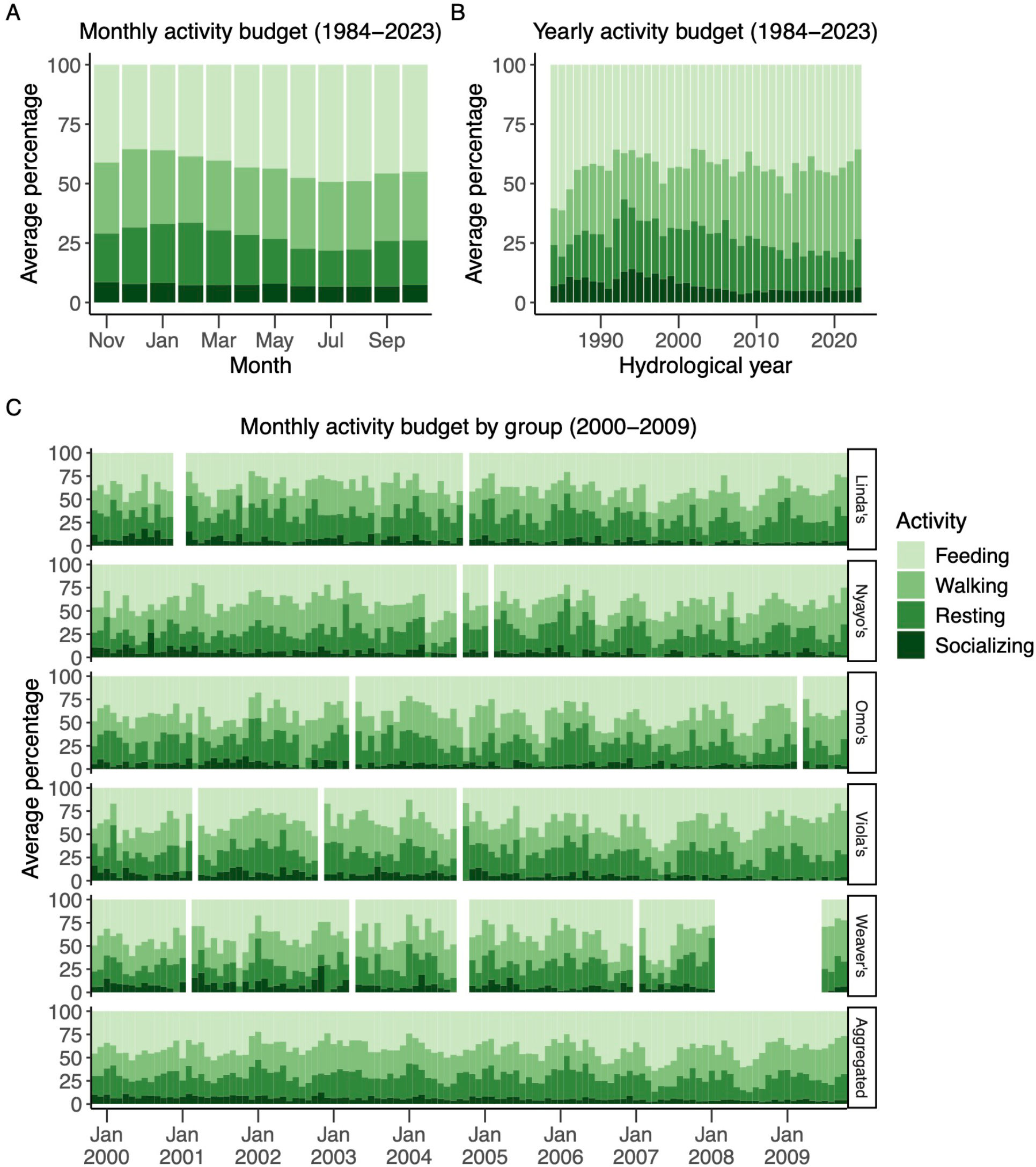
Variation in the percentage of focal sample time that adult females spent feeding, walking, resting, and socializing. (A) in each month, averaged across study groups and years, (B) within each year from 1984-2023, averaged across study groups, and (C) within five social groups, averaged within months from November 1999 to October 2009 (i.e., hydrological years 2000-2009) and averaged within months across the five social groups over the same time period (‘Aggregated’ panel). ‘Socializing’ included time that the focal individual spent grooming, being groomed, and involved in other social interactions with conspecifics. In (C), white spaces indicate group-months in which we believe focal animal sampling was too sparse to produce a statistically representative sample of the activity budgets (n < 10 focal animal samples).

**Figure 4.**
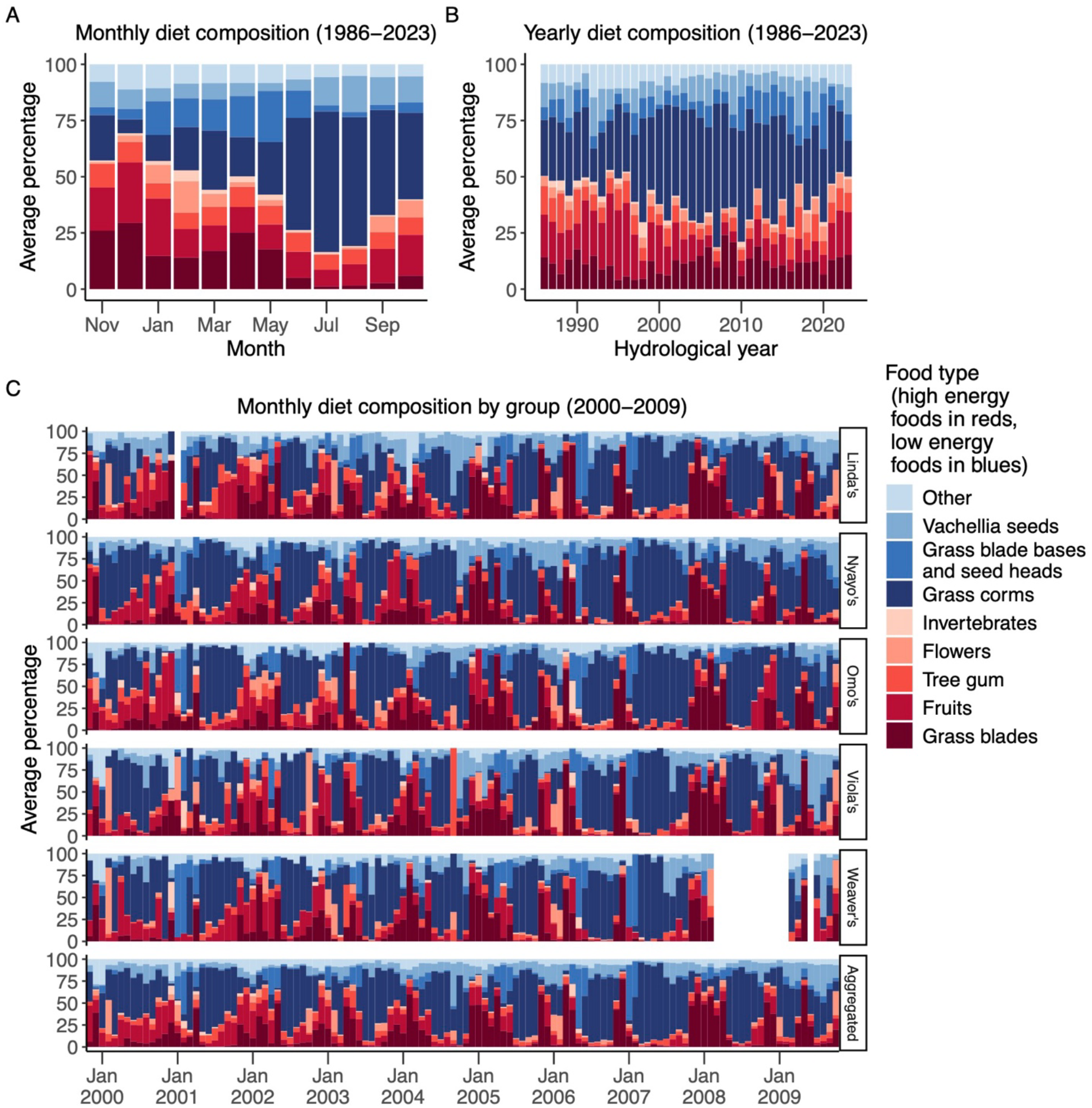
Variation in the percentage of total feeding time spent on each major food type. (A) in each month, averaged across years and across study groups, (B) within each year from 1986-2023, averaged across study groups, and (c) within five study groups, averaged within months from November 1999 to October 2009 (i.e., hydrological years 2000-2009) and averaged within months across the five social groups over the same time period (‘Aggregated’ panel). Presumed high-energy food types are represented in shades of red and low-energy food types are represented in shades of blue. In (C), white spaces indicate group-months in which we believe that focal animal sampling was too sparse to produce a statistically representative sample of diet composition (n < 75 minutes of feeding observed).

These activity budget and diet data were collected from 21 unique social groups across 161-180 group-years and 1,682-1,905 group-months. Because we did not collect activity budget and diet data prior to 1984, rows for those years contain ‘NA.’ After 1984, ‘NA’ values could result from several reasons, including: observations were attempted, but no focal samples contained in-sight observations; focal samples were not collected due to in-progress group fissions or fusions; or other logistical obstacles prevented detailed focal sampling.

### Weather monitoring

The fourth data set published here (weather data) resulted from measurements of daily minimum and maximum temperature and daily rainfall from analog min-max thermometers and rain gauges kept in our field camp. In all years, the thermometer and rain gauge were within a few meters of each other in an area that we term our ‘weather station’. However, the location of the weather station changed over the years, once because we moved our field camp approximately 7.1 km (in 1992), and after that because we moved the weather station to various locations within our field camp as the layout of camp changed (distances between subsequent locations were generally less than 150 m). We include a unique identifier to indicate the weather station where the data were collected and the location (latitude/longitude) of each station. These data were recorded from 1976 to 2022 for temperature and 1976 to 2023 for rainfall. From these daily measurements, we calculated for each month and year: i) the average minimum daily temperature, ii) the average maximum daily temperature, and iii) the total volume of rainfall.

### Data Records

The data are accessible at **link**. Descriptions for each column in each data record are included in table_descriptions.xlsx.

#### Social group history

Aggregated metadata for all ABRP study groups observed for any length of time. File social_group_history.csv (21 social groups) provides information on time periods of observation and social group fissions and fusions for each study group. These metadata are essential for interpreting the group-level data summaries below.

#### Weather station locations

Latitudes and longitudes for each of the 7 weather stations from which weather data were collected between 1976 and 2023 in file weather_station_locations.csv. Exact coordinates were not recorded for stations BC2 and BC3.

#### Monthly data for each study group

Study group composition, activity budget, and diet data are aggregated by month and social group in file monthly_grp_comp_activity_diet.csv (2,372 total group-months). Columns A-C include the social group name, year, and month of data presented in a given row.

##### Study group composition data

(2,372 group-months of data) are included in columns D-I. These data present total group size, numbers of resident adult females, adult males, and immatures, whether the group was undergoing a fission or fusion, and any group-level notes on demography for a social group during a given month from 1971 to 2023.

##### Adult female activity budget data

(1,901 group-months of data) are included in columns J-U. These data present adult female activity budgets (i.e., activity budgets aggregated across all adult females in the group) for each study group during each month and were collected from focal animal samples on adult females between 1984 and 2023. Data were aggregated by pooling all data points collected at one-minute intervals during focal samples, across all adult females in the group, resulting in a group-level estimate of the proportion of total sampling time that the pooled set of adult females spent in each activity (or on each food type).

##### Immature animal activity budget data

(1,682 group-months of data) are included in columns V-AI. These data present immature activity budgets (i.e., activity budgets aggregated across all immature individuals in the group, as described above for adult females) for each study group during each month and were collected from focal animal samples on immatures between 1984 and 2023.

##### Diet data

(1,905 group-months of data) are included in columns AJ-BD. These data present the proportion of feeding time that adult females and immatures spent on different food types for each study group during each month and were collected from focal animal samples on adult females and immatures between 1984 and 2023. Prior to September 1985, observers did not distinguish between different food types when collecting focal animal samples.

#### Monthly data aggregated across all groups

Activity budget and diet data are aggregated across all study groups and weather data are aggregated by month in the file monthly_activity_diet_weather.csv (569 total months). Here, data were aggregated by pooling all data points collected at one-minute intervals during focal samples, across all adult females in all study groups, resulting in a population-level estimate of the proportion of total sampling time that the pooled set of adult females in all study groups spent in each activity (or on each food type). Columns A-B include the year and month of data summarized in a given row. In three of the months shown in the table, data were collected from two weather stations, and each station has a separate row: June 1997, when BC1 was transitioning to BC2; February 2004, when BC3 was transitioning to BC4; and June 2021, when BC4 was transitioning to BC5.

#### Adult female activity budget data

(475 months of data) are included in columns C-N. These data summarize adult female activity budgets during a given month, aggregated across all study groups, and were collected from focal animal samples on adult females between 1984 and 2023.

##### Immature animal activity budget data

(376 months of data) are included in columns O-AB. These data summarize immature activity budgets during a given month, aggregated across all study groups, and were collected from focal animal samples on immatures between 1984 and 2023.

##### Diet data

(478 months of data) are included in columns AC-AW. These data present the proportion of feeding time that adult females and immatures spent on different food types during a given month, aggregated across all groups, and were collected from focal animal samples on adult females and immatures between 1984 and 2023. Prior to September 1985, observers did not distinguish between different food types when collecting focal animal samples.

##### Weather data

(569 months of data) are included in columns AX-BA. These data include the average daily temperature minimum, temperature maximum, and total rainfall for each month and were collected from a series of different weather stations between 1976 and 2023.

#### Yearly data for each study group

These data parallel the monthly data for each study group described above but are aggregated at the level of the hydrological year; in this case, data were aggregated by pooling all data points collected in each study group during the entire hydrological years, resulting in an annual group-level estimate of the proportion of total sampling time that the pooled set of adult females spent in each activity (or on each food type). Social group composition, activity budget, and diet data are aggregated by hydrological year and study group in file hydroyear_grp_comp_activity_diet.csv (216 total group-years). Columns A-B include the study group name and hydrological year of data presented in a given row.

##### Study group composition data

(216 group-years of data) are included in columns C-H. These data present total group size, numbers of resident adult females, adult males, and immatures, whether the group was undergoing a fission or fusion, and any group-level demography notes for a social group during a given hydrological year from 1971 to 2023.

##### Adult female activity budget data

(180 group-years of data) are included in columns I-T. These data present adult female activity budgets for a study group during a given hydrological year and were collected from focal animal samples on adult females between 1984 and 2023.

##### Immature animal activity budget data

(161 group-years of data) are included in columns U-AH. These data present immature activity budgets for a study group during a given hydrological year and were collected from focal animal samples on immatures between 1984 and 2023.

##### Diet data

(180 group-years of data) are included in columns AI-BC. These data summarize adult female and immature diets for a study group during a given hydrological year and were collected from focal animal samples on adult females and immatures between 1984 and 2023. Prior to September 1985, observers did not distinguish between different food types when collecting focal animal samples. All diet data for hydrological year 1985 were categorized as ‘Unclassified’ in this sheet as distinct food types were only collected for in September and October.

#### Yearly weather data

Weather data are aggregated by hydrological year in file hydroyearly_weather_data.csv (47 years and 54 rows of data). These data include average daily temperature minimum and average daily temperature maximum assessed over each hydrological year, and total rainfall for each hydrological year. They were collected from a series of different weather stations between 1976 and 2023.

### Technical Validation

We strive to validate the data we collect by implementing various precautions, safety checks, and database management practices. Detailed explanations of these procedures are available online^24,25^. In the field, skilled observers recorded focal animal sample data as handwritten records between January 1984 and 1999, and electronically beginning in August 1999 (with Psion Workabout units through July 2015 and thereafter with the Prim8 Android app on Samsung tablets^29^). Focal data were sent to database managers in the United States monthly between 1984 and approximately 2005, and on a weekly basis from 2005 until 2015; focal data recorded since 2015 have been sent daily. Handwritten data, including census and weather data, were proofread in the field every Saturday by multiple field team members, who searched for potential inconsistencies or incomplete data. At the end of each month, all handwritten data were proofread again, and (since 2014) scanned and emailed to database managers in the United States; before 2014, handwritten data were mailed to the United States each month. Additionally, on the last four observation days of each month, the field observers collected a reduced number of focal samples in order to collect other types of data, including wounds and pathologies and signs of physical maturation in males. They also used this time to ensure that different observers were reliably identifying the same individuals, which is particularly important for maturing infants and juveniles who can experience rapid changes in coat color and size.

Upon the arrival of the data in the United States, database managers sorted and proofread the data again, discussing any potential errors with the field team. Data were ultimately uploaded to our PostgreSQL database, Babase^30,31^. Babase has built-in validation checks that prevent the uploading of several types of erroneous or impossible data points. However, data were typically first uploaded to a test database with the same validation checks so that any remaining errors were corrected prior to uploading data to Babase. The source code for Babase is freely available and downloadable^25^.

### Usage Notes

The social group history file provides metadata that are necessary to understand how social groups have changed over time via fissions and fusions and should be referenced when analyzing group-level data.

The data on activity budgets and diet were recorded during a limited number of hours each day: focal animal samples were typically only conducted between 0700 and 1100 or between 1400 and 1800, although the daily time periods for focal animal sampling have varied somewhat over the past 40 years. This means that our activity data are representative of baboon behavior during these time periods. In addition, when conditions were not favorable to focal animal sampling (e.g., the group was walking quickly or spread over a very wide area) we missed many more point samples (i.e., recordings of behavior on the minute, within focal samples) than when the animals were resting or close to each other, and these missed point samples may result in bias in which activities were recorded in some circumstances. For instance, in this population the amount of focal data recorded by observers is negatively correlated with the proportion of time individuals spend walking. This correlation is likely caused by frequent movement making it more difficult for observers to maintain visual contact with focal individuals, leading to more out of sight focals or interruptions in sampling. These potential sources of bias in the animals’ activities and diet composition should be taken into consideration when interpreting the activity budget data.

The data on rainfall and temperature were collected from different weather stations, which reflect changes in the devices used to collect the data or the location of the station. It is likely that the weather readings reflect some systematic variation as a function of weather station identity. We include a unique identifier for the weather station that collected each weather data record published here. Temperature data collected by one weather station (ID: BC6) in 2022 and 2023 are not published here because during this period, the station’s analog min-max thermometer was replaced with a digital minimum-maximum thermometer that produced anomalously high readings. In addition to variation as a function of weather station, the study groups have ranged over a wide and shifting area and we suspect that the study area is characterized by spatial heterogeneity in precipitation at fine temporal scales. This spatial heterogeneity means that our rainfall data provide a coarse but not a fine-grained measure of rainfall experienced by the study animals. For example, on a given day, it is possible that some rain fell within the home range of one study group, but not within the home range of another study group.

## Code Availability

R^33^ code used to generate figures is available at **link** and utilizes R packages *tidyverse, ggpubr, data*.*table, cowplot, ggarchery* and *ggnewscale*^33–38^.

## Acknowledgments

We thank Jeanne Altmann for her instrumental role in stewarding the Amboseli Baboon Research Project and in designing many of the ABRP’s long-term data collection protocols. We gratefully acknowledge the support of the National Science Foundation and the National Institutes of Health for the majority of the data represented here, currently through R01AG071684, R01AG075914, and R61AG078470. Current support for field-based data collection also comes from the Max Planck Institute for Evolutionary Anthropology, and we thank Duke University, Princeton University, and the University of Notre Dame for financial and logistical support. In Kenya, our research was approved by the Wildlife Research Training Institute (WRTI), Kenya Wildlife Service (KWS), the National Commission for Science,

Technology, and Innovation (NACOSTI), and the National Environmental Management Authority (NEMA). We also thank the University of Nairobi, the Institute of Primate Research (IPR), the National Museums of Kenya, the members of the Amboseli-Longido pastoralist communities, the Enduimet Wildlife Management Area, Ker & Downey Safaris, Air Kenya, and Safarilink for their cooperation and assistance in the field. We are particularly thankful to the ABRP long-term field team (R.S. Mututua, S. Sayialel, J.K. Warutere, I.L. Siodi, and L. Musembei) and to T. Wango and V. Oudu for their assistance in Nairobi. Design and programming for the ABRP database, Babase, is provided by K. Pinc. This research was approved by the IACUC at Duke University, University of Notre Dame, and Princeton University, and the Ethics Council of the Max Planck Society and adhered to all laws and guidelines of Kenya. A complete set of acknowledgments of funding sources, logistical assistance, and data collection and management is available at http://amboselibaboons.nd.edu/acknowledgements/.

